# Multi-plane Imaging of Neural Activity From the Mammalian Brain Using a Fast-switching Liquid Crystal Spatial Light Modulator

**DOI:** 10.1101/506618

**Authors:** Rui Liu, Neil Ball, James Brockill, Leonard Kuan, Daniel Millman, Cassandra White, Arielle Leon, Derric Williams, Shig Nishiwaki, Saskia de Vries, Josh Larkin, David Sullivan, Cliff Slaughterbeck, Colin Farrell, Peter Saggau

## Abstract

We report a novel two-photon fluorescence microscope based on a fast-switching liquid crystal spatial light modulator and a pair of galvo-resonant scanners for large-scale recording of neural activity from the mammalian brain. The utilized imaging technique is capable of monitoring large populations of neurons spread across different layers of the neocortex in awake and behaving mice. During each imaging session, all visual stimulus driven somatic activity could be recorded in the same behavior state. We observed heterogeneous response to different types of visual stimuli from ~ 3,300 excitatory neurons reaching from layer II/III to V of the striate cortex.

## 1. Introduction

Recording neural activity from a large population of neurons in the living brain with physiologically relevant sampling rates and cell-type specificity is particularly important for developing a mechanistic insight into brain function and its disorders [1,2]. The advent of genetically encoded neural activity indicators [3], transgenic reporter mouse lines expressing those indicators in specific cell types [4], and modern fluorescence microscopy, have made optical recording of neural activity a promising approach towards this ambitious goal.

Despite intense efforts, large-scale recording from the mammalian brain of awake and behaving animals remains a challenge for the following reasons. First, most neural circuits extend over considerable space, requiring a method capable of sampling neural activity in a large volume of brain tissue [5]. Second, mammalian brain tissue is opaque due to its strong scattering profile, making it hard to image neurons deep in the brain [5]. Lastly, suitable imaging methods must overcome motions induced by awake and behaving animals.

The mammalian neocortex is horizontally organized as six morphologically distinct layers, I-VI, with different roles in information processing. Cortical columns, composed of narrow chains of interconnected neurons of different types extending vertically through cortical layers, are perceived as basic modular units of the neocortex [6]. Multi-plane imaging is a conceptually straightforward approach to achieve large-scale recording of neural activity by utilizing the laminar structure of neocortical columns. In particular, more evidence has accumulated over the years showing that locomotion and pupil dilation are accompanied by multiple effects on sensory processing [7]. Changes of behavior state from one imaging session to another make it challenging for experiments designed to understand how object representations are transformed between layers of a cortical column using conventional single-plane two-photon microscopy. Therefore, multi-plane imaging has become essential to record neural activity at different cortical layers in the same behavior state. In addition, when studying the functional connectomics of the neocortex, this approach also facilitates a co-registration of imaging data from the same neocortical column obtained by functional optical imaging and large-scale electron microscopy [8,9], because both methods sample the brain tissue uniformly along spatial dimensions.

Following this strategy, new imaging techniques based on electrically tunable lenses have demonstrated their potential to record neural activity from two different cortical layers [10]. However, the utility of these devices for reliable routine imaging is often limited by their optical performance, including intrinsic aberrations, sensitivity to environmental conditions, slow switching speed, and high manufacturing tolerances. The remote focusing approach moves a low-inertia mirror at the focal plane of a secondary, remote focusing objective and can achieve fast focusing. But it is prone to potential laser damage at the focal spot on the remote focusing mirror, especially when high laser powers are used for deep imaging [11, 12]. Another approach, combining a deformable mirror and the concept of temporal focusing, is able to record in a continuous volume, remains limited by relatively small imaging volumes (i.e., 50 × 80 × 20 μm with 5 axial planes) [13] and only in weakly scattering specimens. With other advanced optical recording technologies, such as random-access scanning with acousto-optical deflectors [14] or sculpted light microscopy [15], overcoming behavior related motions is difficult. Furthermore, sculpted light microscopy also requires custom-designed ultrafast lasers to be fully functional [15].

On the other hand, overdrive techniques recently applied to liquid crystal spatial light modulators (SLM) [16, 17], have reduced their switching time to 3 ms [18, 19], making them compelling devices for fast-focusing multi-plane imaging. Furthermore, precise phase control of a large number of pixels (i.e., ~ 250,000 on a single SLM) allows us to generate the high-fidelity wavefront patterns needed to focus over a wide axial range.

In this work, we report a novel two-photon fluorescence microscope based on an overdrive liquid crystal SLM, resonant-galvo scanners, and a conventional ultrafast laser, capable of recording neocortical activity of thousands of neurons spread across different cortical layers in the brain of an awake and behaving mouse. During each imaging session, we recorded all visual stimulus driven somatic activity in the same behavior state.

Compared to earlier reports concerning the use of SLM for fast volume imaging [18, 20], there are several advances in this work. First, Yang et al [18] used an SLM to split a single laser beam into multiple beamlets while we employ the SLM to rapidly switch between different focal planes. We believe our approach is advantageous in the sense that we don’t sacrifice two-photon excitation efficiency by splitting the peak power among focal planes. To maintain the same signal to noise ratio as the one in conventional two-photon microscopy, Yang et al [18] has to increase the applied laser power with the number of beamlets, resulting in a 4~9 times stronger out-of-focus fluorescence background for all imaging planes when imaging a densely labeled cortical column. Second, different from Yang et al [18, 20], the resonant galvo scanner scheme we used in this study results in a larger field of view (FOV) (i.e., 500 × 500 *μ*m *v.s*. 250 × 250 *μ*m) for calcium imaging and thus avoids the stitch of multiple small FOVs within a focal plane. Furthermore, the improved frame rate for multi-plane imaging offered by the resonant galvo scanner in our imaging scheme helps recording neural activity from a larger field of view (i.e., 500 × 500 *μ*m) without experiencing much in-plane non-rigid motions. Lastly, beyond a proof-of-concept study [20], we applied our imaging method to functional imaging of neural activity from thousands of neurons in the primary visual cortex of awake and behaving mice, revealing an interesting response behavior of certain neurons to various types of visual stimuli. These results demonstrate the utility of our imaging method for meaningful biological studies.

## 2. Methods

### 2.1. Microscope Design and Operation

The microscope design is shown in Fig.1. A laser beam from an ultrafast Ti:Sapphire oscillator (Spectra-Physics, Insight X3) with a wavelength of 940 nm and a pulse duration of ~ 120 fs was expanded by a factor of 6 to fill the active area of the SLM (Meadowlark Optics, HSP-512). The SLM was conjugated to the resonant scanner by a 4f lens system composed of L3 (Thorlabs, AC254-100-B) and L4 (Thorlabs, AC254-040-B). The 12 kHz X resonant scanner (Cambridge Technology, CRS 12K) was conjugated to the Y galvo scanner (Cambridge Technology, 6215H) by two identical scan lenses L5 and L6 (Thorlabs, SL50-3P) also arranged in a 4f configuration. Similarly, the y galvo was conjugated to the back focal plane of the objective lens (Nikon, N16XLWD-PF) with a 4× Keplerian telescope, containing L7 (Thorlabs, SL50-3P) and L8 (Thorlabs, TL200-3P). The objective was mounted on a piezo actuator (PZT, Physical Instruments, P-725KHDS). The back pupil of the objective was under-filled to achieve an effective excitation numerical aperture (NA) of 0.45. The prechirping configuration for the ultrafast laser pulses was optimized to achieve the highest excitation efficiency. Emitted fluorescence was collected by the same objective as used for excitation and reflected by a dichroic mirror (Semrock, F705-Di01-25×36) towards the active area of a photomultiplier (Hamamatsu, H10770PA-40 SEL).

**Fig. 1.**
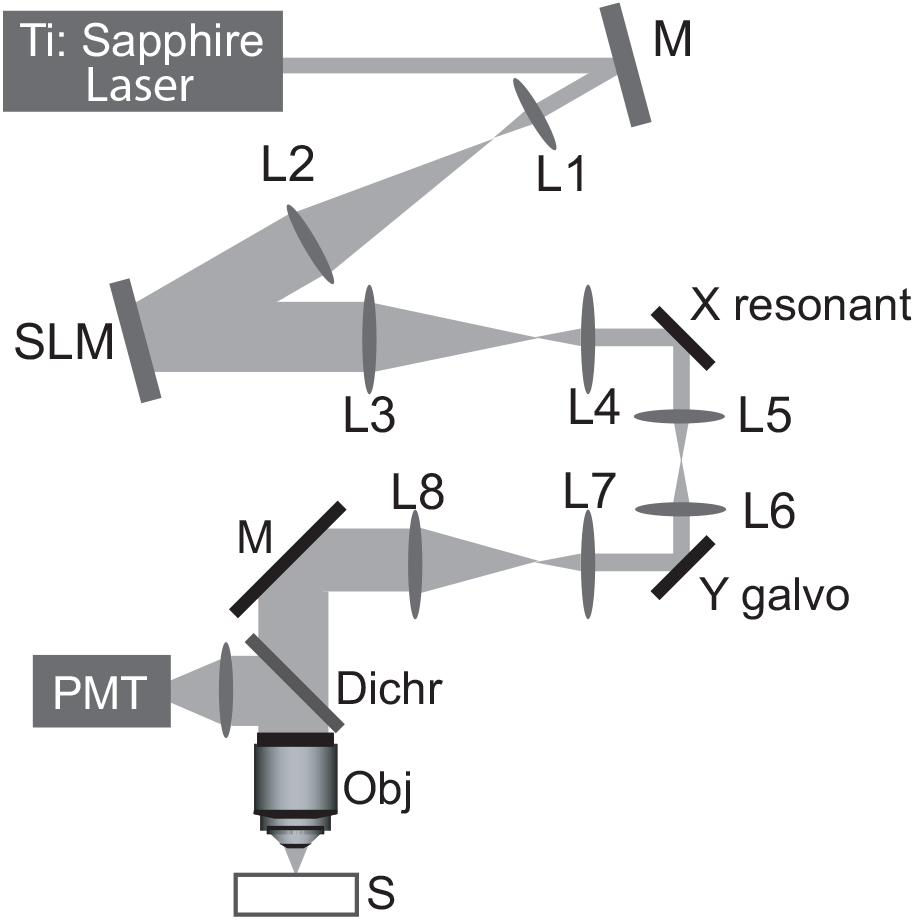
Schematic diagram of our microscope system. M, mirror; L1–L8, lens; SLM: liquid crystal spatial light modulator; Dichr: dichroic mirror; Obj: microscope objective; PMT: photomultiplier; S: Sample; X resonant: resonant scanner in x direction; Y galvo: galvo scanner in y direction

Our microscope is based on the modular *in vivo* multiphoton microscopy system (MIMMS) design developed at HHMI Janelia Research Campus [21] but includes significant modifications, as shown in Fig.2. To maintain the strict conjugation between optics in the microscope, we could not move the objective as it is typically used in the MIMMS. Instead, we used a 3D translational stage (Dover, XYR-8080 and ZE-100) to load and unload the animal. The movable microscope head (Sutter Instruments) was rotated along the optical axis of L8 by 23.7 deg and the raised optical breadboard, containing most of the mounted microscope optics, was rotated by another 18 degrees with respect to the animal. This geometrical arrangement ensures that during the experiments, the optical axis of the microscope objective is aligned perpendicular to the surface of the cranial window, with a tolerance of less than 1 deg. Eye-tracking and behavior-monitoring cameras were used to track pupil movements and whole body movements of the animal during imaging sessions, respectively. The X galvo in Fig.2 was not used in this study and kept stationary.

**Fig. 2.**
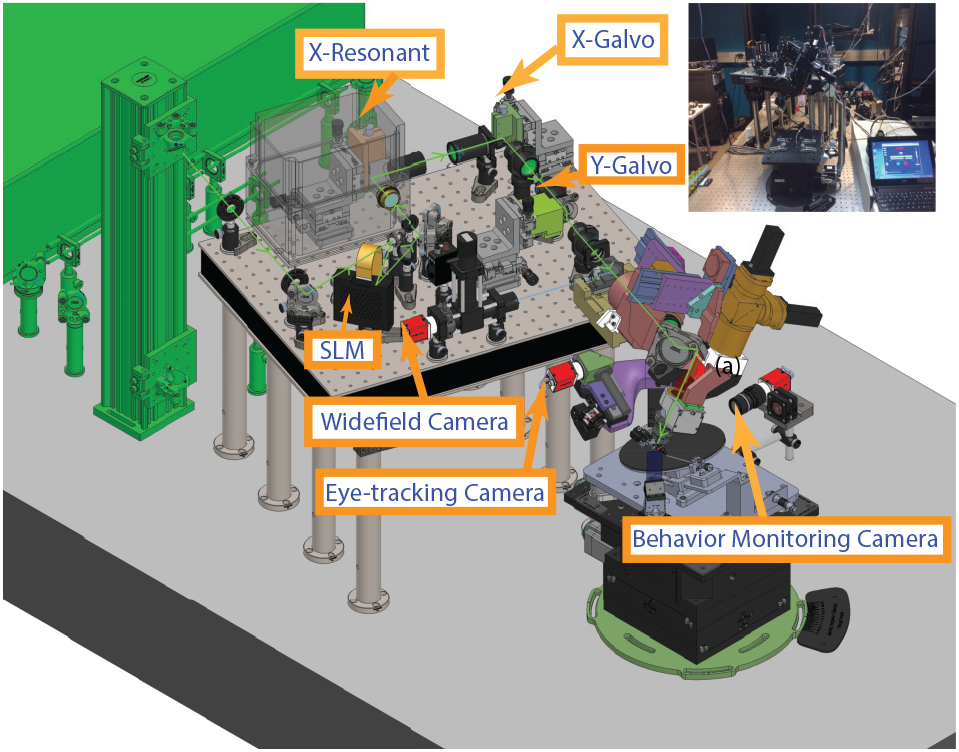
3D CAD model for the microscope design. Inset: picture of our experimental setup

Due to the strict conjugation between SLM and back focal plane of the objective, any wavefront changes at the SLM are equivalent to a direct manipulation at the Fourier plane of this objective. Therefore, changing the collimation of the laser beam at the back pupil of the objective could be directly controlled by “displaying” defocus wavefront patterns with different magnitudes on the SLM. As a result, the microscope can focus at different axial planes by applying different defocus wavefront patterns. Before each imaging session, adaptive optics procedures were performed on 2 *μ*m fluorescent beads to correct for residual optical aberrations due to possible slight misalignments using a procedure described in [22], ensuring we are applying ideal defocus wavefront patterns in the system.

Updating the defocus patterns on the SLM was synchronized to a single-frame resonant-galvo scan for image acquisition by a handshake protocol. Single-frame image acquisition based on resonant-galvo scanning was implemented with ScanImage 2016 (Vidrio Technologies), while the SLM was controlled and synchronized with a custom software written in LabVIEW 2015 (National Instruments). After updating the SLM pattern, image acquisition of each frame was hardware-triggered. Upon completing a frame, the falling edge of the frame clock was utilized to update the SLM pattern for the next frame. By cycling through a limited number of different defocus patterns, the system sequentially and repeatedly acquires images at different depths in the mouse brain.

### 2.2. Surgery

Dexamethasone (3.2 mg/kg, S.C.) was administered to the mouse three hours prior to surgery. Carprofen (5-10 mg/kg, S.C.) was then applied before the incision procedure to remove the skin on top of the head. Under isoflurane anesthesia (5% initially, and 1.5-2 % during the surgery), a custom designed headframe was attached to the skull using Metabond (Parkell) and a 5 mm craniotomy was performed at center coordinates 2.8 mm lateral, 1.3 mm anterior to lambda. The craniotomy was sealed with a stack of three #1 glass coverslips (two 5 mm and one 7 mm glued together) and Vetbond. Metabond cement was further applied to secure the cranial window. The animal’s health was monitored for 7 days post-surgery, i.e., overall status (alert and responsive), cranial window clarity, and brain health.

### 2.3. Visual Stimulation

Visual stimuli were presented to the right eye of the mouse using an ASUS PA248Q LCD monitor with 1920 × 1200 pixels. The center of the monitor was positioned 15 cm away from the mouse eye, covering a visual space of 120°× 95°. The monitor was gamma calibrated using a USB-650 Red Tide Spectrometer (Ocean Optics) and had a mean luminance of 50 *cd/m*^2^, measured with a SpectroCAL MKII Spectroradiometer (Cambridge Research Systems). All custom visual stimulation scripts were written using PsychoPy [23, 24] and Python 2.7.

Visual stimuli included drifting gratings, natural movies, and a display of mean luminance grey to record spontaneous activity, with a total length of 62 minutes. The timing sequence of those visual stimuli was arranged as described in Fig. 3(d). The drifting gratings stimulus consisted of a full field drifting sinusoidal grating at a single spatial frequency (0.04 cycles/degree). It was presented to the mouse eye at 8 different directions (separated by 45 degrees) and 5 temporal frequencies (1, 2, 4, 8, 15 Hz). Each grating was presented for 2 seconds with 1 second of mean luminance grey between trials. Each grating condition was presented 15 times, in a randomized order, with 30 blank sweep trials interleaved. Natural movie 1 and 2 clips were two different movie clips extracted from the opening scene of the movie *Touch of Evil*. The length of the natural movie 1 is 2 min, while the natural movie 2 is 30 sec in length. Each movie clip was presented 10 times. A spherical warping was applied to all types of the visual stimuli to compensate for the close viewing angle of the mouse, ensuring size, speed, and spatial frequency were constant across the monitor as seen by the mouse.

**Fig. 3.**
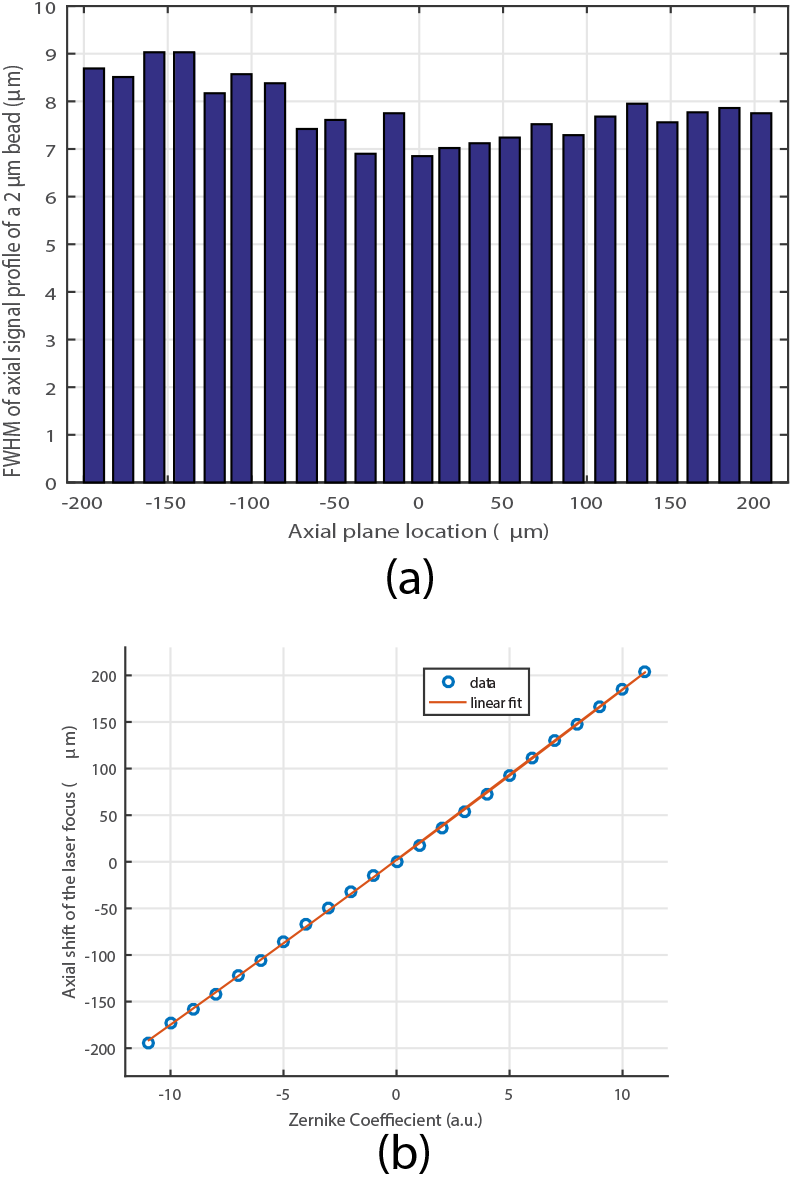
(a) FWHM of the axial profile of z stack images of 2 *μ*m fluorescent beads does not degrade significantly over an axial range of 400 *μ*ms (b) Axial shifts of focal planes from its nominal position are practically linear with the Zernike defocus coefficients

### 2.4. Imaging Workflow

Each mouse to be imaged on our system first underwent intrinsic signal imaging (ISI) to obtain a retinotopic map to define visual area boundaries and ensure two-photon calcium imaging could be targeted to consistent retinotopic locations. ISI time series were acquired under light anesthesia with 1-1.4% isoflurane administered using somnosuite (Kent Scientific, model #715) to measure the hemodynamic response to the visual stimulus. Sign maps were obtained from the averaged ISI time series from a minimum of 30 sweeps of drifting gratings in each direction. The resulting ISI map was automatically processed, resulting in a segmented and annotated map to guide later two-photon imaging sessions.

After a successful ISI imaging session, each mouse went through a two-week period of habituation to make sure it was habituated to head fixation, visual stimulation, and two-photon imaging.

During two-photon imaging sessions, the mice were positioned on a rotating disk, free to run at will during the experiments. A magnetic shaft encoder (US Digital) attached to this disk recorded the running speed at 60 samples per second. Pupil dilation and the animal posture were monitored and recorded using two separate industrial cameras (Allied Vision, Mako G-032B). Two-photon image acquisition, visual stimulation, eye-tracking, running speed recording, and mouse body monitoring are all synchronized by means of a custom software written in Python 2.7.

More details regarding surgery, imaging procedures, and visual stimulation have been previously reported in [25–27]. All experiments and procedures were approved by the Allen Institute Animal Care and Use Committee.

## 3. Results

### 3.1. System Characterizations

One potential concern with this refocusing method is that it is not operating the objective the way it was designed for (i.e. under the assumption that the rays are entering the entrance pupil in a collimated manner from different angles). Therefore, additional spherical aberrations are unavoidable from the perspective of aberration theory [28]. For this reason, we characterized the performance of our microscope by taking z stack images of 2 *μ*m fluorescent beads using a PZT to move the objective lens while the laser was focused to different axial planes by means of defocus patterns displayed on the SLM. Fig.3(a) demonstrates that the FWHM of axial images of 2 *μ*m beads did not degrade significantly over an axial range of 400 *μ*m when using an effective excitation NA of 0.45, consistent with other previous report [18]. However, we anticipate a decrease of the acceptable axial range of the point spread function (PSF) degradation when using a higher effective NA. Therefore, to maintain the optical performance of the system over an extended axial range in this scenario, adaptive optics procedures have to be employed to correct for those defocus induced aberrations [29, 30]. By replacing lens L3 with one of different focal lengths, the overall beam expansion ratio from the SLM to the back focal plane of the objective can be adjusted to achieve different effective excitation NAs. As shown in Fig.4, with an effective NA of 0.65 but without adaptive optics, the PSF degraded quickly when the SLM was used to focus tens of microns away from the nominal focal plane. When adaptive optics procedures as described in [22] were applied on top of different defocus wavefront patterns, the PSFs at different depths were recovered over an axial range of ~ 100 *μ*m.

**Fig. 4.**
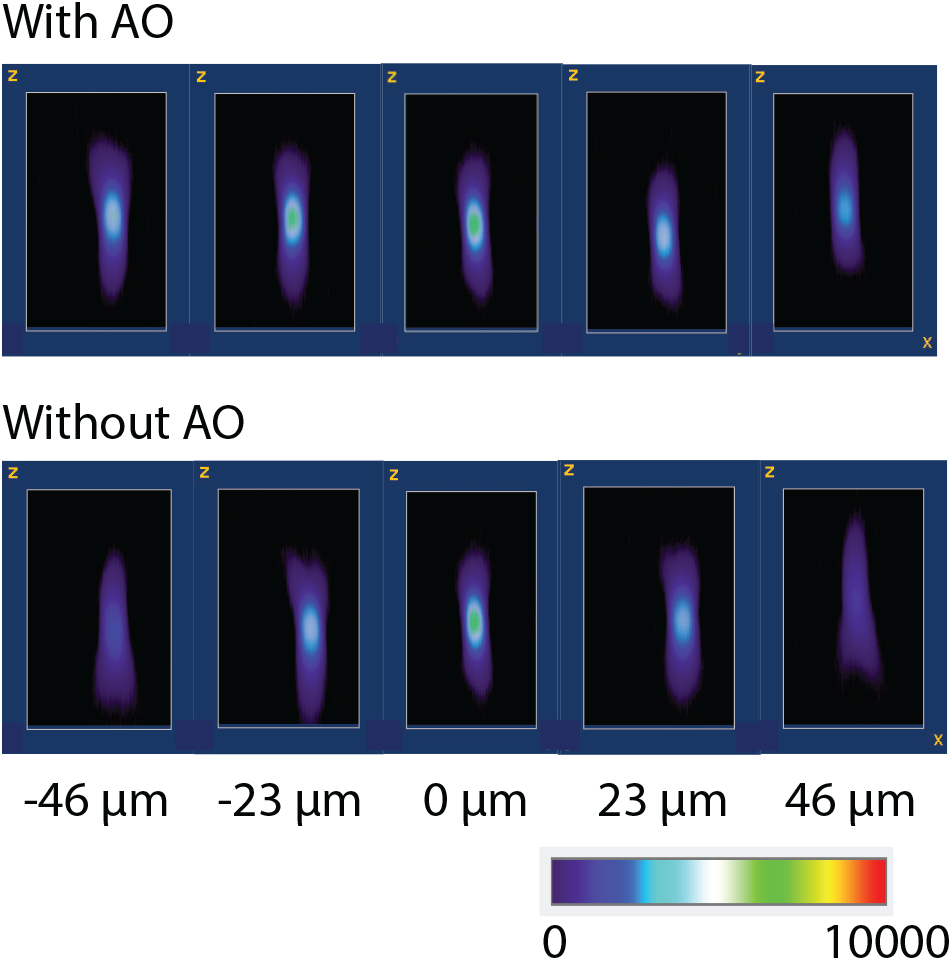
Axial profile of z stack images of 2 *μ*m fluorescent beads with different axial shifts using the SLM defocusing mechanism with an effective numerical aperture of 0.65. Upper: with adaptive optics; Bottom: without adaptive optics

We also characterized the relationship between the Zernike defocus term coefficients and the actual axial shifts of the focal plane. Our measurements show that the locations of actual focal planes obtained with different defocus patterns are almost linearly proportional to the coefficients of the Zernike defocus term. The fit shown in Fig.3(b) yields *y* = 0.034*x*^2^ + 17.9753*x* + 1.7056. This finding agrees well with a previous report on implementing a similar refocusing concept using deformable mirrors [31]. We utilized this practically linear dependence to design different multi-plane imaging configurations with focal planes arbitrarily separated within the axial range specified in Fig.3(a).

### 3.2. *Functional Optical Imaging from Awake and Behaving Mice* in vivo

Our microscope is able to focus at different planes without mechanical inertia, within an axial range of 400 *μ*m. Employing an overdrive liquid crystal SLM minimized the switching time between frames. The exact imaging depths and the separations between different image planes can be extracted from the linear dependence shown in Fig.3(b). In this work, we demonstrated three different configurations for multi-plane imaging using a fast-switching SLM. As shown in Fig.5(a), we are able to image 6 axial planes spread from layer II/III to layer V with moderate sampling rate (6.5 Hz). Fig.5(b) shows another configuration in which two imaging planes were located at two different cortical layers but with a higher sampling rate of 19.5 Hz. By arranging the imaging planes 23 *μ*m between each other, we could image a tissue volume of (500 × 500 × 92 *μ*m) at ~ 7.8 Hz.

**Fig. 5.**
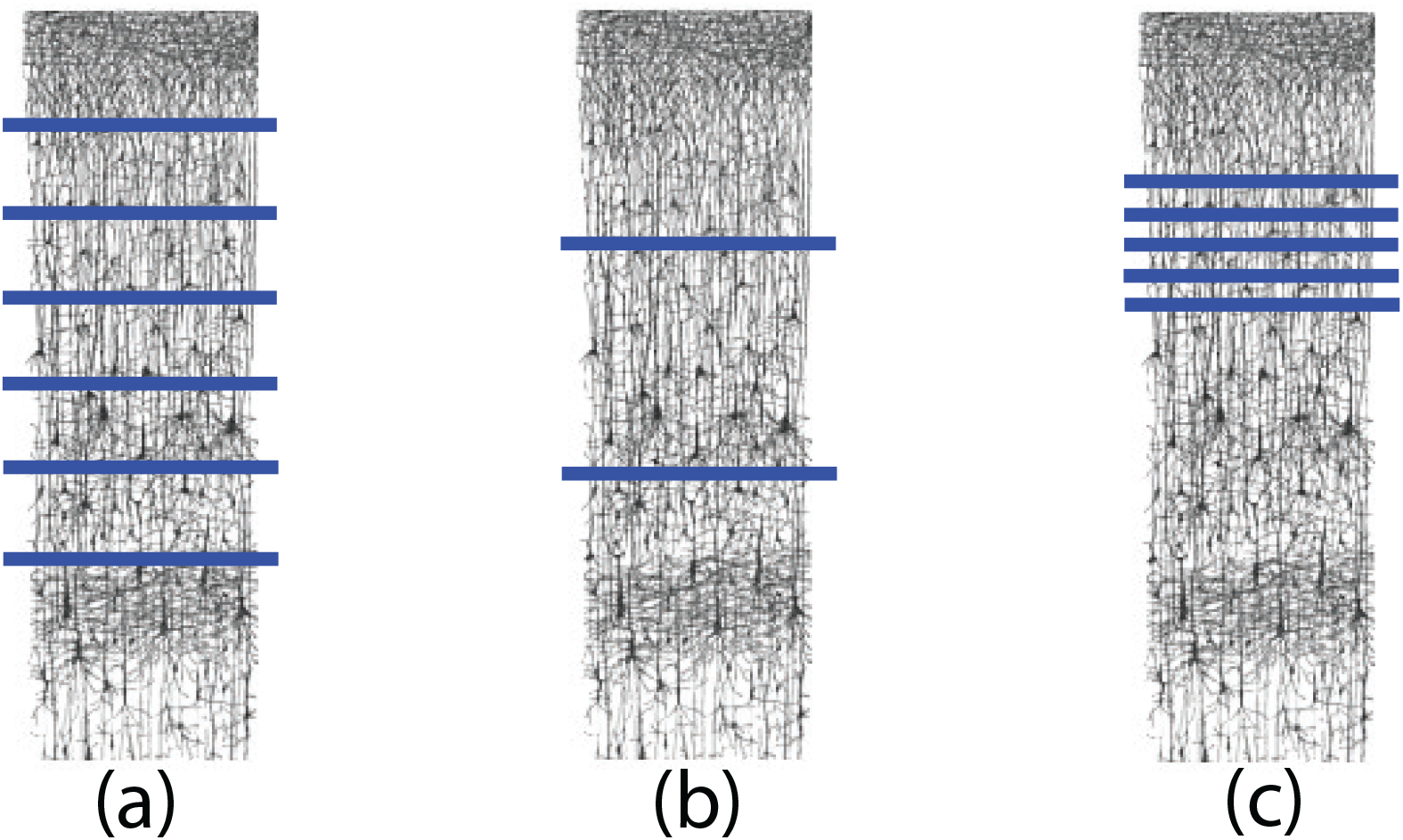
(a) Multi-plane imaging of 6 planes separated by 50 *μ*m each; (b) Multiplane imaging of two axial planes separated by 204 *μ*m (c) Volumetric imaging of 5 imaging planes separated by 23 *μ*m each. The neural circuit diagram is adapted from [7]

We imaged neural activity from neurons in layer II to V of an awake and behaving transgenic mouse (Slc17a7-IRES2-Cre;Camk2a-Tta;Ai94 (TITL-GCaMP6s) line), visiting 6 different focal planes, each plane with afield of view (FOV) of 500 *μ*m × 500 *μ*m(512 × 512 pixels) separated by 50 *μ*m as shown in Fig.5(a). Calcium activity movies were recorded together with synchronized visual stimulus, running speed and pupil dilation. Calcium imaging data for all imaging planes were processed by Suite2P for motion correction, segmentation, and neuropil removal [32]. Fig.6(a) shows the maximum intensity projections of all images after motion correction at focal depths of 500 *μ*m, 450 *μ*m, 400 *μ*m, 350 *μ*m, 300 *μ*m, and 250 *μ*m, respectively. The frame rate for each plane was about 6.5 Hz. Fig.5(b) shows the ROI segmentation at different imaging depths. Distinct segmentation patterns at different planes clearly demonstrated that those images originated from different depths of the brain. In addition, we observed the number of large cells is gradually increasing, starting from an imaging depth of 400 *μ*m. At an imaging depth of 500 *μ*m, most of the segmented neurons are larger than those segmented in shallow layers, which agrees well with the anatomy of mouse neocortex. The related calcium imaging movie can be found in Visualization 1. In the densely labeled cortical column of the slc17a7-IRES2-Cre;Camk2a-Tta;Ai94 (TITL-GCaMP6s) mouse line, we also observed that the out-of-focus fluorescence background got increased with greater imaging depth. However, the degraded signal to background ratio did not prevent us from performing further image processing, such as the motion corrections and segmentation, down to ~ 550 *μ*m (data not shown here). Throughout all 6 imaged focal planes, we are able to extract neural activity from a total of ~3,300 cells at a sampling rate of ~ 6.5 Hz in a single imaging session.

**Fig. 6.**
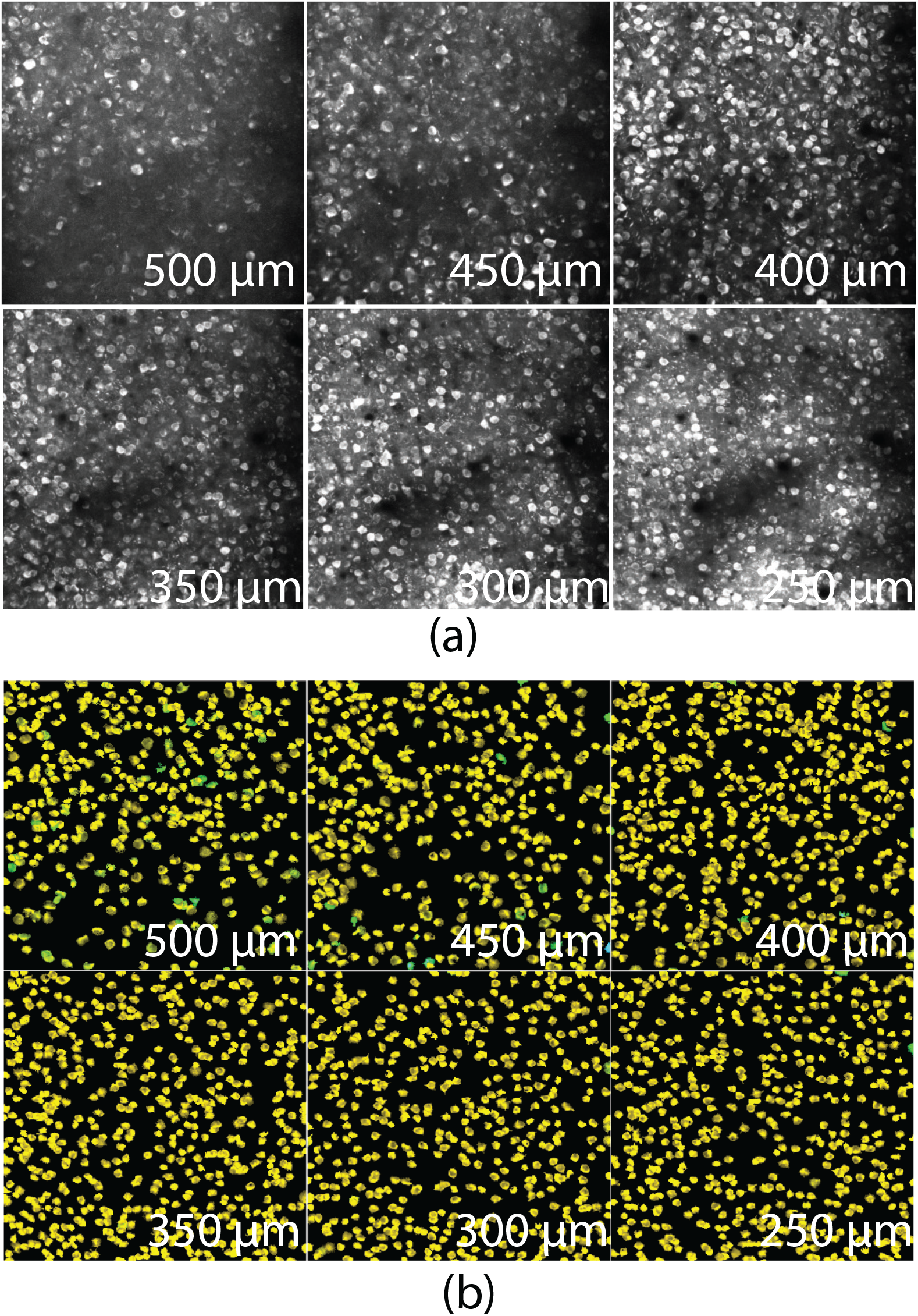
(a) Maximum intensity projection of the calcium imaging movie at different depths (see Visualization 1); (b) Segmented somata identified by Suite2P at different depths

To characterize visual responses of neurons in the mouse visual cortex, a combination of stimuli was used, including drifting gratings (30 min) and 2 natural movies, movie 1 (20 min), movie 2 (5 min), and a 5 minute grey screen to record spontaneous activity. Fig.7(b) shows activity traces from a selected group of neurons defined in Fig.7(a). The time sequence of the combined visual stimuli is shown in Fig.7(c), exactly aligned to the neural activity traces in Fig.7(b). During the one-hour imaging session, neurons 5 and 6 responded mainly to drifting gratings, while neurons 3 and 4 responded mainly to natural movies. Many neurons behave like neurons 1 and 2, responding to different types of visual stimuli. All this demonstrated a heterogeneous response of neurons in the primary visual cortex to the same visual stimulus and locomotive state. Fig.7(d) presents the recorded running speed, showing the mouse was running during most of the imaging session. We further analyzed the tuning curve of the 4 neurons responding to drifting gratings in Fig.7(b). Fig.8 shows very similar tuning curves for neurons 5 and 6, while neuron 1 only responded to one direction of drift and neuron 2 was less tuned towards any particular direction. This observation is consistent across different cortical layers. The orientation selectivity index (OSI) for all segmented somata were calculated for each imaging plane. As shown in Fig.9, we observed many cells with the classical tuning behaviors at different cortical layers. However, a significant portion of excitatory neurons are responding to visual stimuli but don’t show a preference towards any particular tuning orientation, which suggests even excitatory neurons in the primary visual cortex may have complicated cortical processing functionality beyond edge detection.

**Fig. 7.**
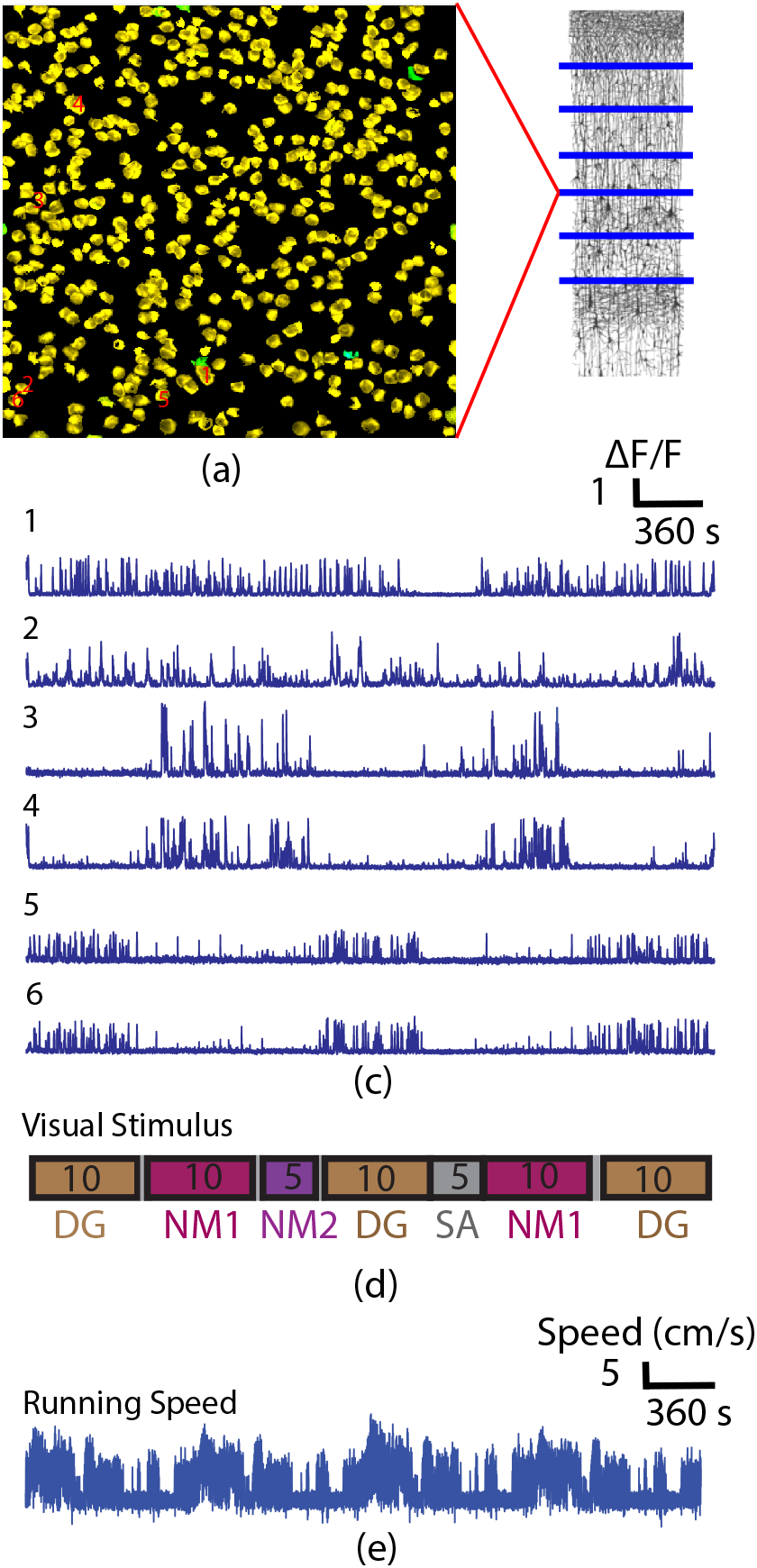
(a) Illustration of selected ROIs in the imaging plane at a depth of 400 *μ*m; (b) Neural activity traces extracted from (a); (c) Visual stimulus composed of drifting grating (DG), natural movie 1 (NM1), natural movie 2 (NM2), and grey screen for recording spontaneous activity; (d) Running speed recorded during the imaging session. The neural circuit diagram in (a) is adapted from [7]

**Fig. 8.**
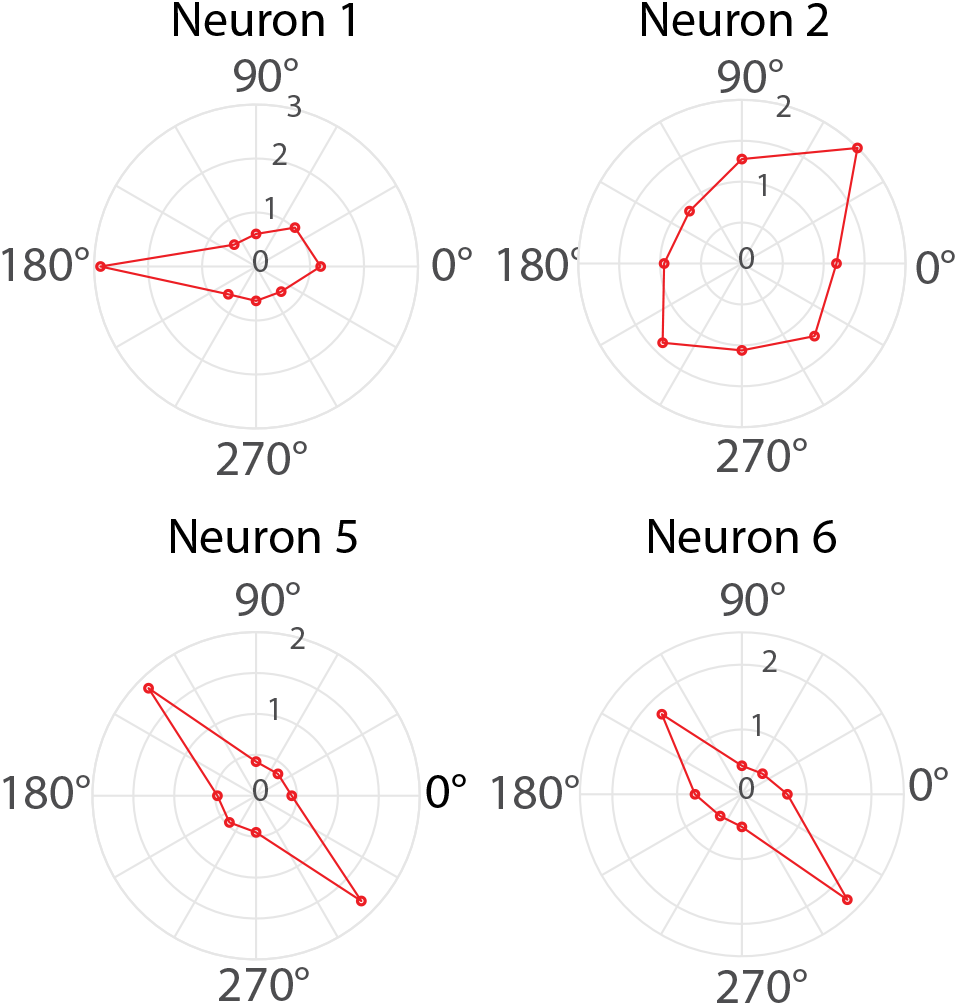
Tuning curves of selected neurons, the radial magnitude is in units of ΔF/F

**Fig. 9.**
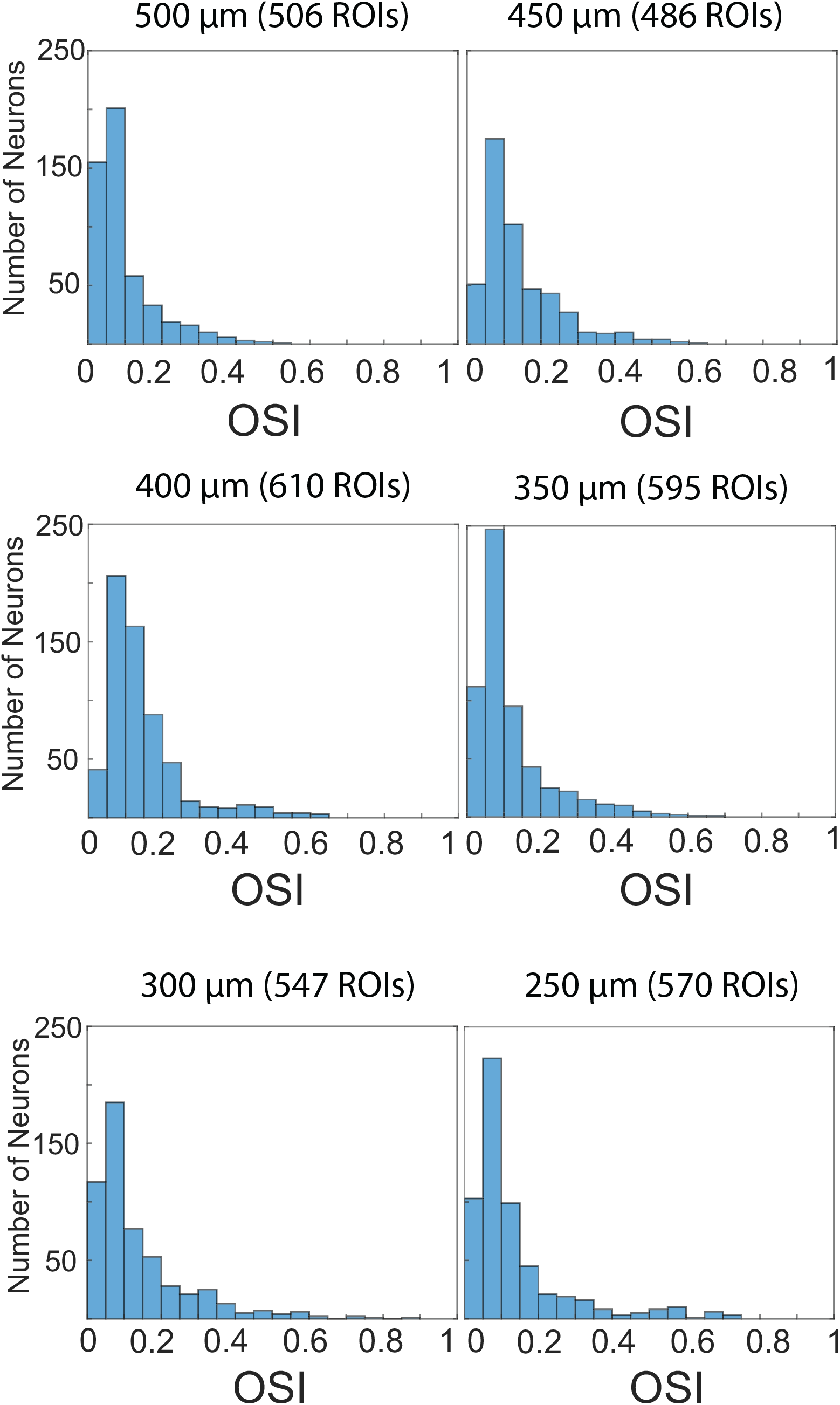
Histogram of the orientation selectivity index for all segmented somata at different imaging depths

## 4. Discussions

Compared to [18], our imaging method has to compromise between the frame rate and the number of imaging planes with a given image acquisition time for each plane as shown in Equation. 1

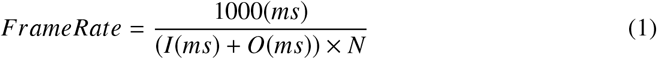

Where I is image acquisition time for each plane, O is overhead switching time, including SLM switching and software overhead, and N is number of imaging planes. For example, for a 12 kHz resonant scanner operated in bidirectional mode and largest possible FOV (i.e., 512×512 pixels), the frame rate would be about 6.5 Hz for a total of 6 imaging planes, with 22.4 ms for image acquisition and 3 ms for overhead switching. The frame/volume rate can be increased by reducing the total number of imaging planes or reducing the number of resonant scan lines for each frame.

The total number of recorded neurons is limited by the labeling density of the neurons within the sampling volume as well as the sampling rate used in the imaging session. With about 2.5 Hz sampling rate in [32], we could image 16 planes and sample about ~ 8800 neurons in total. Our current imaging field of view (500 × 500 *μ*m) is limited by the expansion ratio defined by L7 and L8 in Fig.1. With the improved optical design for L7 and L8, we could further increase the field of view for each imaging plane and thus enable us to record more neurons from each imaging session.

Different from [32], our imaging planes are aligned perfectly along the laminar cortical layers and are not required to be arranged to be close to each other. Using the imaging configuration illustrated in Fig.5(b), Fig. 10 shows maximum intensity projections of the calcium imaging movie at two focal planes located at layers II/III and V of a transgenic mouse (Rorb-IRES2-Cre;Camk2a-tTA;Ai94(TITL-GCaMP6s)), respectively, with a separation of 204 *μ*m and identical frame sizes. Here, frame rates for both planes are about 19.5 Hz. By separating the imaging planes 23 *μ*m from each other, we could also image a volume of (500 × 500 × 92 *μ*m) at ~ 7.8 Hz. Fig. 11 shows maximum intensity projections for images from 5 closely spaced axial planes. One thing worth noting, when imaging planes were arranged close to each other, higher NA (NA=0.65) is used. Adaptive optics can tighten the PSF at different axial focal planes and help to minimize signal crosstalk between planes. However, it seemed hard to completely remove all contaminations simply because of the heterogeneous sizes of neurons.

**Fig. 10.**
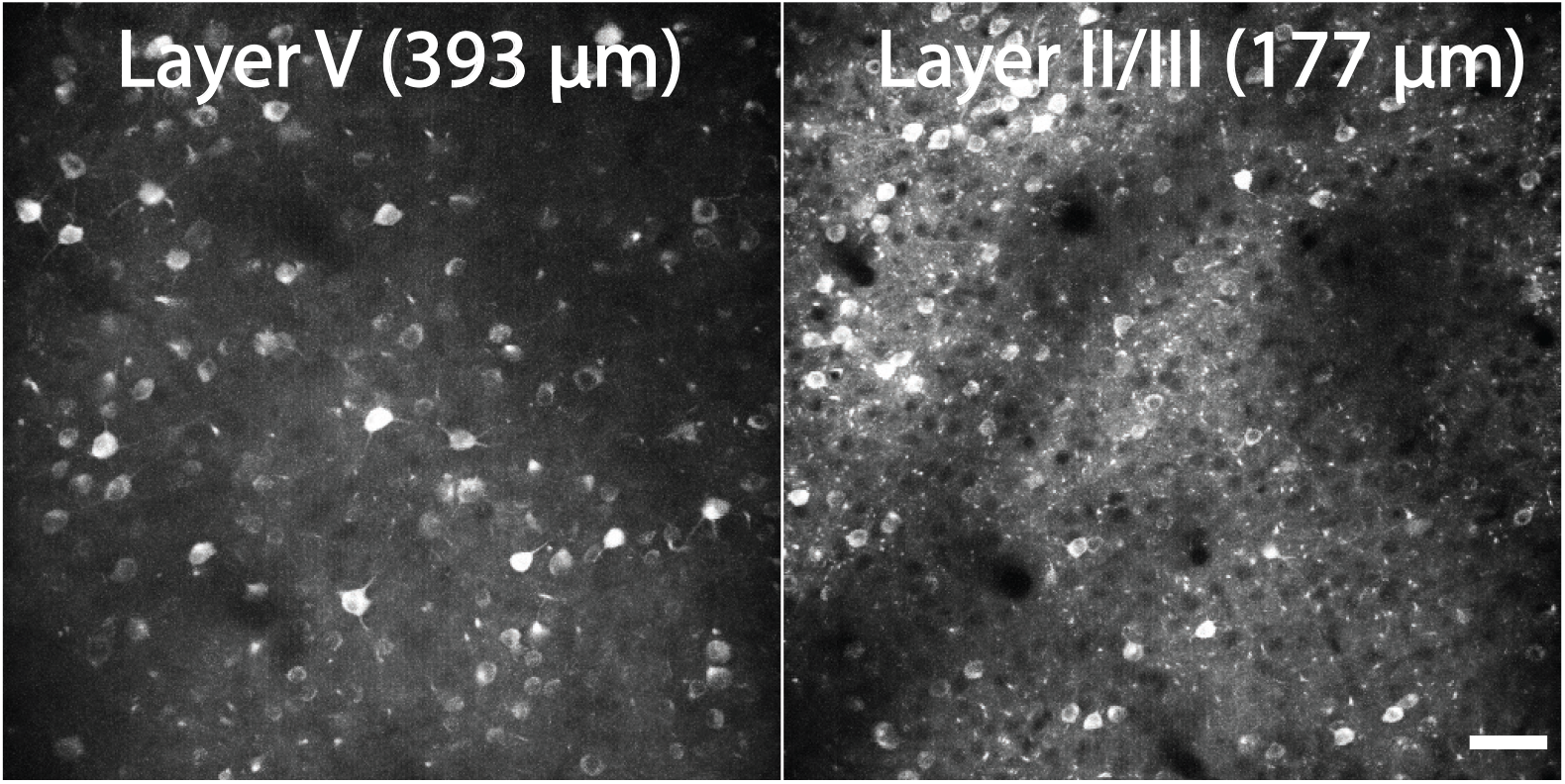
Maximum intensity projection of calcium imaging movies at two different axial planes with 204 *μ*m separation. (see Visualization 2). Scale bar: 50 *μ*m

**Fig. 11.**
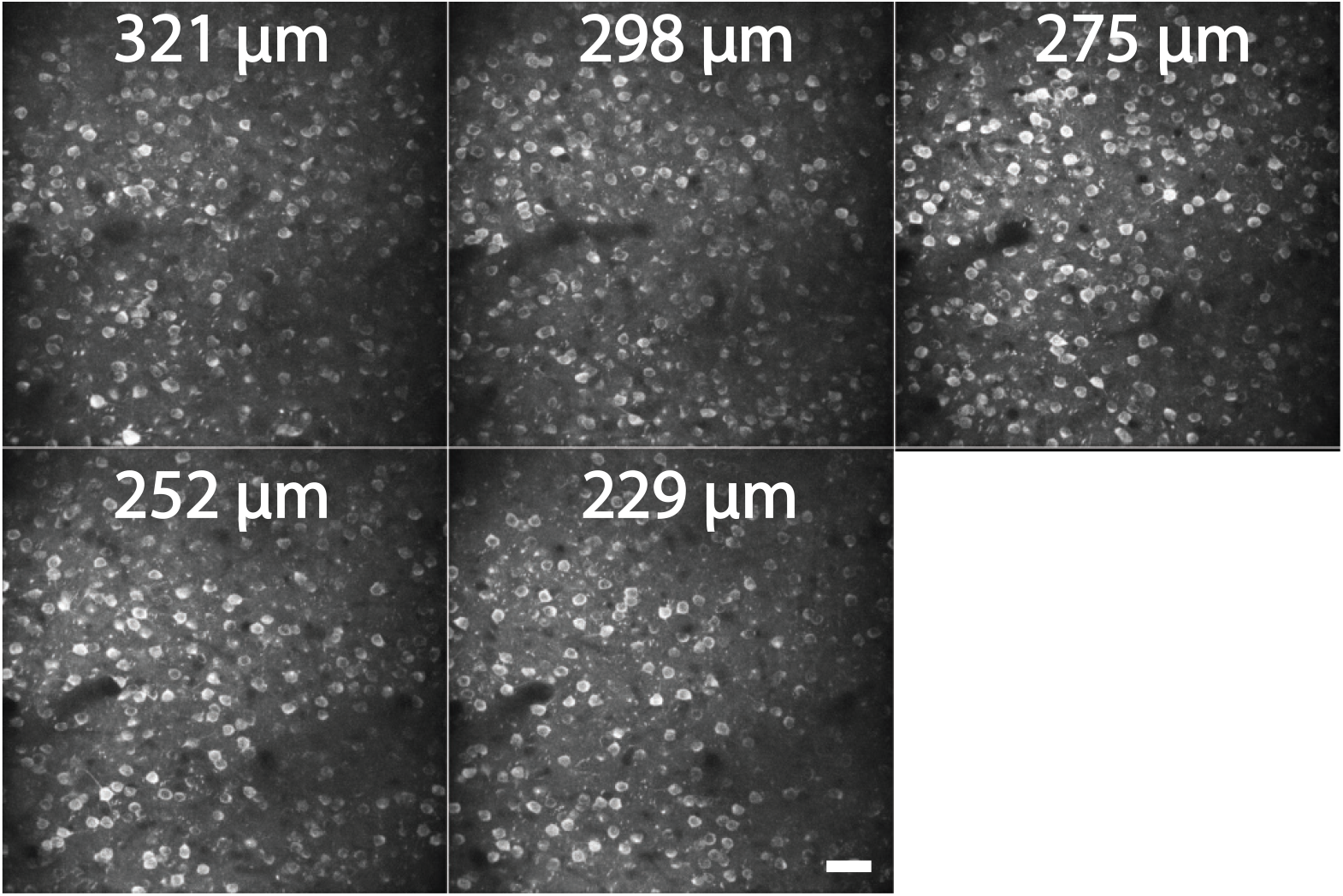
Maximum intensity projection of calcium imaging movies at five different axial planes with 23 *μ*m separation. (see Visualization 3). Scale bar: 50 *μ*m

In summary, we have demonstrated multi-plane imaging of neural activity using a fast-switching liquid crystal spatial light modulator, capable of recording neocortical activity of thousands of neurons spread across different cortical layers of the brain of awake and behaving mice. During each imaging session, all visual stimulus driven somatic activity could be recorded in the same behavior state. This imaging scheme is readily configurable for volumetric imaging with focal planes arranged close to each other, e.g., 23 *μ*m separation between planes. Similarly, it can also be reconfigured to record neural activity from two cortical layers separated far apart by a few hundred microns. In addition, we demonstrated the utility of this method by performing multi-plane imaging across different cortical layers to record heterogeneous response from neurons to different types of visual stimuli. This method does not have special requirements for fluorescence labeling, such as the labeling sparsity [33] and multi-color labeling. The extended PSF due to the low numerical aperture could be beneficial for increasing the number of neurons recorded in each imaging session utilizing the increasingly sophisticated demixing algorithms [32, 34]. It would also be interesting to combine this approach with the multi-foci approach reported in [18] to further improve the overall throughput in sparsely labeled specimen. With the ongoing improvements of fast-switching liquid crystal spatial light modulator in terms of switching speed and pixel density [19], we anticipate the frame rate for multi-plane imaging to be ultimately limited by the raster-scan frame acquisition time. For this reason, replacement of resonant scanners with even faster polygon mirrors [35] will be the next step to further improve the acquisition rate and recording capacity for multi-plane imaging. Furthermore, temporal multiplexing [36] can be combined with our multi-plane imaging approach to boost the imaging throughput within the fluorescence lifetime limit of available neural activity indicators. Lastly, it is also possible to utilize our multi-plane imaging scheme to dynamically correct for the licking-induced motions during the behavior experiment as shown in [37].

## Acknowledgments

The authors would like to thank many staff members of the Allen Institute, especially Hongkui Zeng for GCaMP6 reporter lines, and the In Vivo Sciences team for performing the surgeries.

